# Conserved mycobacteriophage protein uses structural mimicry to displace host initiation factors

**DOI:** 10.64898/2026.10.05.756390

**Authors:** David R. Parker, Shrestha Ghosh, Adela Dujsikova, Paul Dominic B. Olinares, Sarah Narrowe Danielsson, Grisna Isabel Prensa, Doyeon Kim, Florian Andreas Rosenberger, Arne Elofsson, Matthias Mann, Elizabeth Campbell, Andrea Fossati

**Author notes:** Contributing authors. These authors contributed equally to this work (David R. Parker, Shrestha Ghosh, Adela Dujsikova).

## Abstract

Compact viral genomes encode sophisticated strategies to reprogram host gene expression, often by co-opting rather than replacing RNA polymerase. How bac-teriophages achieve such control in Mycobacteria remains largely unknown. Here, time-resolved interaction proteomics of mycobacteriophage D29 infection iden-tifies a conserved two-protein module, gp53-gp52, that binds the host RNA polymerase. Affinity purification confirmed association of both proteins with the polymerase, while native mass spectrometry demonstrated that gp53 displaces the host sigma factor. Structural modelling places gp53 at the sigma-binding surface of RNA polymerase, consistent with molecular mimicry of sigma-factor engagement. The module is conserved throughout Cluster A mycobacteriophages and occurs in fused or split genomic architectures. These findings suggest sigma-factor displacement as a mechanism of transcriptional takeover in viruses of mycobacteria and show how a compact viral module can repurpose an essential host molecular machine.

## Introduction

Bacterial viruses (bacteriophages, or phages) provide powerful systems for probing and exploiting bacterial physiology, as they have evolved highly efficient strategies to reprogram core cellular processes to replicate [1]. Central to successful infection is rapid host takeover, during which phages redirect gene expression away from host genes and toward phage replication [2]. Transcription represents a key control point in this process, and rapid access to and control over host transcriptional resources is therefore critical for productive phage infection [3]. Phages have evolved diverse strate-gies to exploit this vulnerability. These include redirection of host RNA polymerase (RNAP) toward phage promoters, inhibition of host transcriptional programs[3], and the deployment of phage-encoded transcriptional[2, 4, 5]. Some phages bypass host transcription by encoding dedicated RNA polymerases, effectively shifting from host rewiring to transcriptional replacement, often accompanied by broad inhibition of the host RNA polymerase [6, 7]. While these mechanisms are well established in model phages like T4[8], how phages manipulate transcription in other bacteria, such as Mycobacteria, remains significantly less understood. This gap is particularly striking given that mycobacteriophages comprise one of the largest and most diverse known phage groups, with more than 2,400 sequenced genomes and 10,000 isolated phages [9, 10]. Several mycobacteriophage clusters, such as A and K, despite being isolated on fast-growing laboratory hosts (*M. smegmatis*), are able to infect other Mycobac-teria species[11], including slow-growing Mycobacteria such as *M. tuberculosis*[12], highlighting a broad host range across bacteria with vastly different transcriptional profiles, growth rate and life cycles [13]. In both fast- and slow-growing Mycobacteria, transcription is a dominant regulatory bottleneck governed by a single essential RNAP core, comprised of the subunits RpoB, C, a dimer of RpoA and RpoZ (*α*_2_*ββ^′^ω*), is suf-ficient for catalytic activity[14] but requires initiation factors, Sigmas (*σ*), to form the holoenzyme that is capable of recognizing and initiating at promoter sequences [15]. Mycobacterial RNAP requires additional basal factors, including the housekeeping SigA-associated factor, RbpA, for competent initiation[16]. The expanded repertoire of sigma factors further enables extensive transcriptional remodeling in response to stress, growth state, persistence and antibiotic exposure [17–19]. Importantly, despite this host diversity and scale of mycobacteriophage genomes, no phage-encoded multi-subunit RNA polymerase has been identified in sequenced mycobacteriophages to date. This absence suggests that, unlike some model phages, all mycobacteriophages must achieve transcriptional takeover by engaging and modulating the host RNA polymerase rather than disabling and replacing it with a viral encoded polymerase. However, the molecular strategies underlying this process, including whether and which phage-encoded factors directly interact with host RNA polymerase or associated regulators, are unknown. Here, we address this gap using cluster A phage D29, a lytic derivative of L5 phage, as a representative mycobacteriophage model, able to infect both fast- and slow-growing Mycobacteria [20]. By systematically mapping phage-host interactions during infection, we identify two phage proteins (gp53 and gp52) that directly interact with the mycobacteria RNA polymerase. Notably, gene 53 of D29 and its L5 orthologue are annotated in RefSeq as a “sigma-K factor” (NC 001900.2, NP 046868.1; NC 001335.1). To our knowledge this assignment has never been tested experimentally, and it is not reflected in other resources: the same protein is listed in UniProtKB as “Gene 53 protein” at evidence level “predicted” (O64243), and its only domain assignment is to the phage-specific family PF25684 (IPR057899), with no match to any *σ*70-family model. The experiments below provide the first direct test of this annotation. Ectopic expression and subsequent purification in the absence of phage infection recovers them as RNA polymerase associated proteins, while in silico struc-tural analysis suggests similarity to *σ*2 and *σ*3 domains. Comparative genomic analysis reveals broad conservation in A mycobacteriophages, with three genomic architec-tures, where gp53 and gp52 are either fused, partially overlapping or separated. Native mass spectrometry (nMS) assays demonstrates that gp53 binds to Mycobacterial core RNAP and, intriguingly, displaces the housekeeping sigma factor, SigA, from RNAP. Together our data uncover a previously unrecognized mechanism for transcriptional host takeover in mycobacteriophages based on sigma factor displacement.

## Results

### A time-resolved interactome of bacteriophage infection

To investigate how mycobacteriophages remodel host macromolecular assemblies dur-ing infection, we used a co-fractionation mass spectrometry workflow combining size-exclusion chromatography (SEC) with data-independent acquisition (DIA) mass spectrometry (Fig. 1A). Native cell lysates are separated by SEC under non-denaturing conditions, which resolves proteins according to the apparent size of the complexes they belong to rather than their monomeric mass. Each SEC fraction is then analyzed by DIA-MS, enabling consistent, deep quantification of thousands of proteins across the molecular weight range (*>*2 MDa to ∼10 kDa). Proteins that physically associate within the same complex show similar elution profiles, so complexes and infection-induced remodeling can be inferred directly from correlated chromatographic traces using machine learning approaches [21–23]. Because the approach does not rely on affinity purification, it provides an unbiased, systems-level view of endogenous complex organization and how it changes during infection[24]. It captures both stable assem-blies and condition-dependent shifts in complex composition, abundance or apparent size. Here, *M. smegmatis* cultures were infected with phage D29 and sampled at early (30 min) and mid-infection (120 min) alongside uninfected controls. Lysates were fractionated under native conditions into 70 fractions and analyzed by proteomics to identify interactions between host complexes and phage proteins. In total, we iden-tified 5,162 host proteins, 78.2% of the *M. smegmatis* reference proteome (Fig. 1B), including 94.3% of reviewed and 76.4% of unreviewed UniProt entries. Mapping the detected phage proteins onto the D29 genome revealed near-complete coverage of the annotated proteome (72 of 77 ORFs, 93.5%; Fig. 1B). Protein identifications were highly consistent between biological replicates across the fractionation. In fractions 10-65, a median of 81-87% of proteins per fraction were detected in both replicates (Fig. 1C). Overlap was lower only in the earliest fractions, where few proteins elute (on average ∼470 proteins per fraction in fractions 1-8, versus ∼3,170 in fractions 10-65). Elution profiles were also quantitatively reproducible: the median Pearson cor-relation between replicate profiles was 0.88 in uninfected cells and 0.85 and 0.89 at 30 and 120 min post-infection (Fig. 1D). Together, these results support the robustness of the SEC-DIA-MS workflow for large-scale interactome profiling. In every condition, about half of the host proteins showed two or more elution peaks (53.7%, 52.8% and 55.3% in uninfected cells and at 30 and 120 min; Fig. 1E). This is consistent with proteins partitioning between free pools and one or more assemblies, and the stable proportion indicates that infection does not globally change host elution behavior. D29 proteins eluted across the entire SEC range, often in several discrete peaks (Fig. 1F). Finally, SEC-MS recovered well-characterized host complexes, indicating that the approach preserves native interactions in mycobacteria. The subunits of the 30S and 50S ribosomal subunits, core RNA polymerase (RpoA, RpoB, RpoC and RpoZ), the ClpP1/ClpP2 protease and the proteasome (Mpa, PrcA and PrcB) each co-eluted in a tight band (Fig. 1G). The 50S subunit (median apex fraction 15) eluted ahead of the smaller 30S subunit (fraction 23), as expected from their sizes. These data show that co-elution profiles reliably reconstruct endogenous mycobacterial complexes, making the approach suitable for mapping phage-host interactions.

**Fig. 1.**
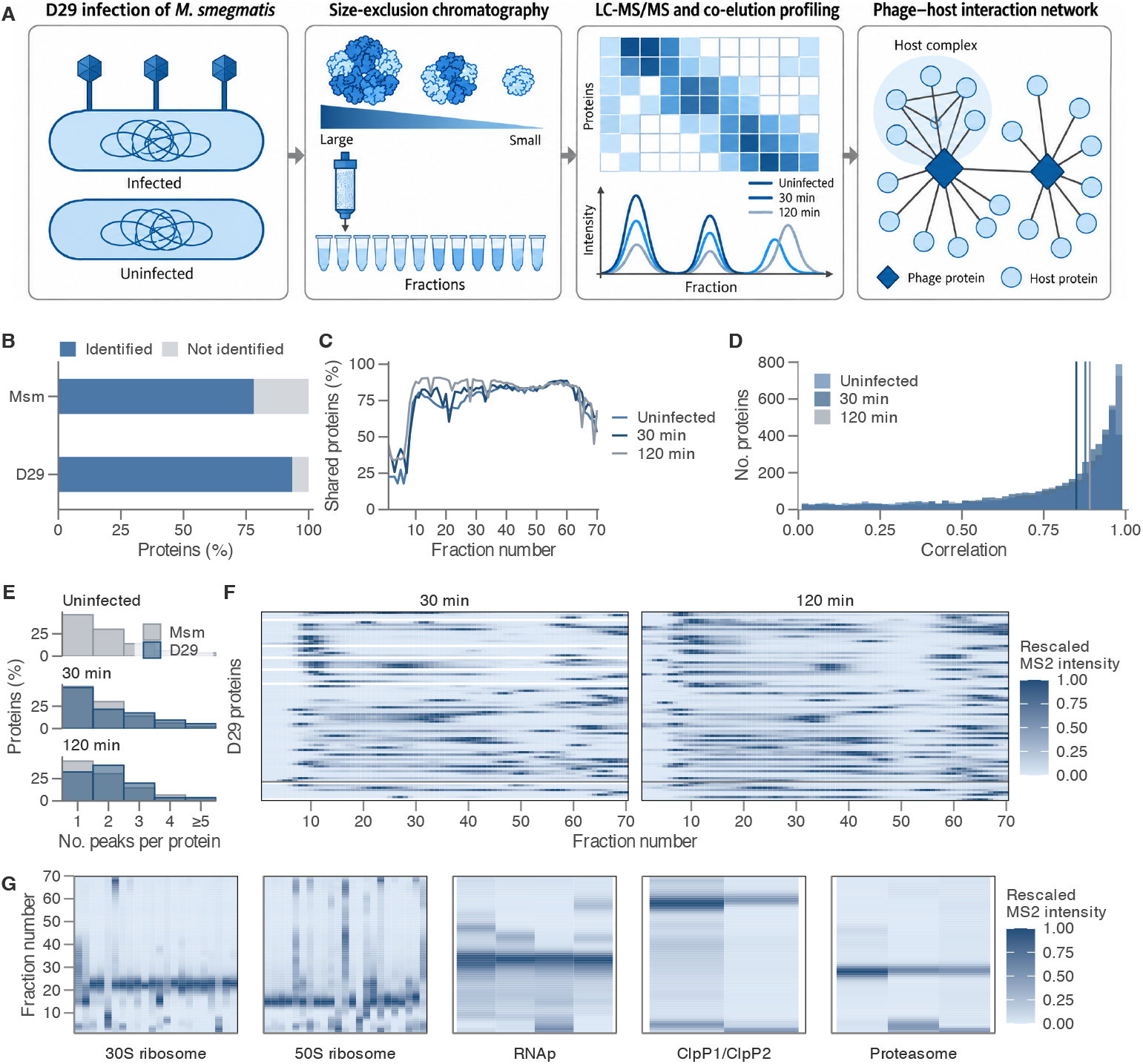
High-throughput interaction proteomics for host–pathogen network mapping during mycobacteriophage D29 infection. **A.** SEC-MS workflow and experimental design. **B.** Proportion of the *M. smegmatis* reference proteome and of annotated D29 ORFs identified (blue) or not (grey). **C.** Percentage of proteins identified in both replicates per SEC fraction, for uninfected cells and 30 and 120 min after infection. **D.** Pearson correlation between replicate elution profiles for each protein; vertical lines mark the condition medians (0.88, 0.85, 0.89). Proteins with *r <* 0 (3–5%) are not shown. **E.** Elution peaks per protein for *M. smegmatis* (grey) and D29 (blue), as a percentage within each group. **F.** Elution profiles of D29 proteins at 30 and 120 min, ordered by gene number (the eight rows below the line lack one). Intensities are replicate-averaged, Gaussian-smoothed (*σ* = 1 fraction) and scaled to 1 per row; white, not detected. **G.** Co-elution of known *M. smegmatis* complexes in uninfected cells. Columns are subunits, the y-axis is fraction number, and colour shows unit-normalized intensity.

### Phage proteins directly interact with the host RNA polymerase

We next asked whether D29 proteins engage essential bacterial macromolecular assem-blies involved in transcription or translation, reasoning that such interactions could reveal mechanisms of host takeover. Across the D29 proteome, gp52 (61 aa), gp53 (234 aa) and gp53.1 (50 aa) showed among the strongest correlations with the elu-tion profile of the mycobacterial RNA polymerase (RNAP) core complex (Fig. 2A). Consistent with this, all three proteins co-eluted with RNAP at both early and mid stages of infection, with closely overlapping chromatographic profiles at 30 and 120 min post-infection (Fig. 2B and Supplementary Fig. S2). Quantification of the RNAP peak fractions recovered the expected *α*_2_*ββ^′^ω* composition of the core enzyme (appar-ent RpoA:RpoB:RpoC:RpoZ ≈ 2.3:1:1.3:0.8), which was unchanged from uninfected cells to 120 min post-infection (Fig. 2C), indicating that the core polymerase is maintained during phage infection. Because the core enzyme is unchanged, any phage-driven change in host transcription would more likely involve which factors load onto the core than the core itself. We therefore looked at sigA, the essential housekeeping sigma factor that directs most transcription in mycobacteria and a common target of phage proteins that redirect host RNAP, such as T4 AsiA. To separate RNAP-bound from free SigA, we compared the amount of SigA in the RNAP peak with that in the SigA monomer peak. The ratio was similar in uninfected cells and at 30 min (1.05 and 0.87) but fell about three-fold by 120 min (0.35) as free SigA accumulated (Fig. 2D), potentially suggesting phage interference with SigA association. To deter-mine whether either protein could associate with RNAP in the absence of additional phage factors, we generated *M. smegmatis* strains expressing Strep-tagged gp52 or gp53 from an acetamide-inducible promoter and performed affinity purification-mass spectrometry (AP-MS)[25, 26] with and without D29 infection. Inducing gp53, but not gp52, slightly decreased growth relative to the empty-vector control, but did not produce a lethal phenotype (Supplementary Fig. S1). Purification of both gp52 and gp53 recovered all four core RNAP subunits, RpoA, RpoB, RpoC and RpoZ, demon-strating that each protein can associate with the host polymerase independently of ongoing phage infection (Fig. 2E,F). Intriguingly, gp52 additionally recovered several sigma factors, including SigA, SigE and SigH, whereas these interactions were not detected with gp53. This difference suggests that gp52 and gp53 engage distinct states or surfaces of the RNAP complex and raise the possibility that gp53 interferes with sigma-factor association. Together, these data identify gp52, gp53 and gp53.1 as D29 proteins whose chromatographic behaviour is closely linked to RNAP during infection and establish gp52 and gp53 as direct RNAP-associated proteins through complemen-tary affinity-purification and biochemical approaches. The presence of gp52 and gp53 in distinct RNAP-containing states, together with their differential recovery of sigma factors, suggested that this conserved phage module may regulate host transcription during infection.

**Fig. 2.**
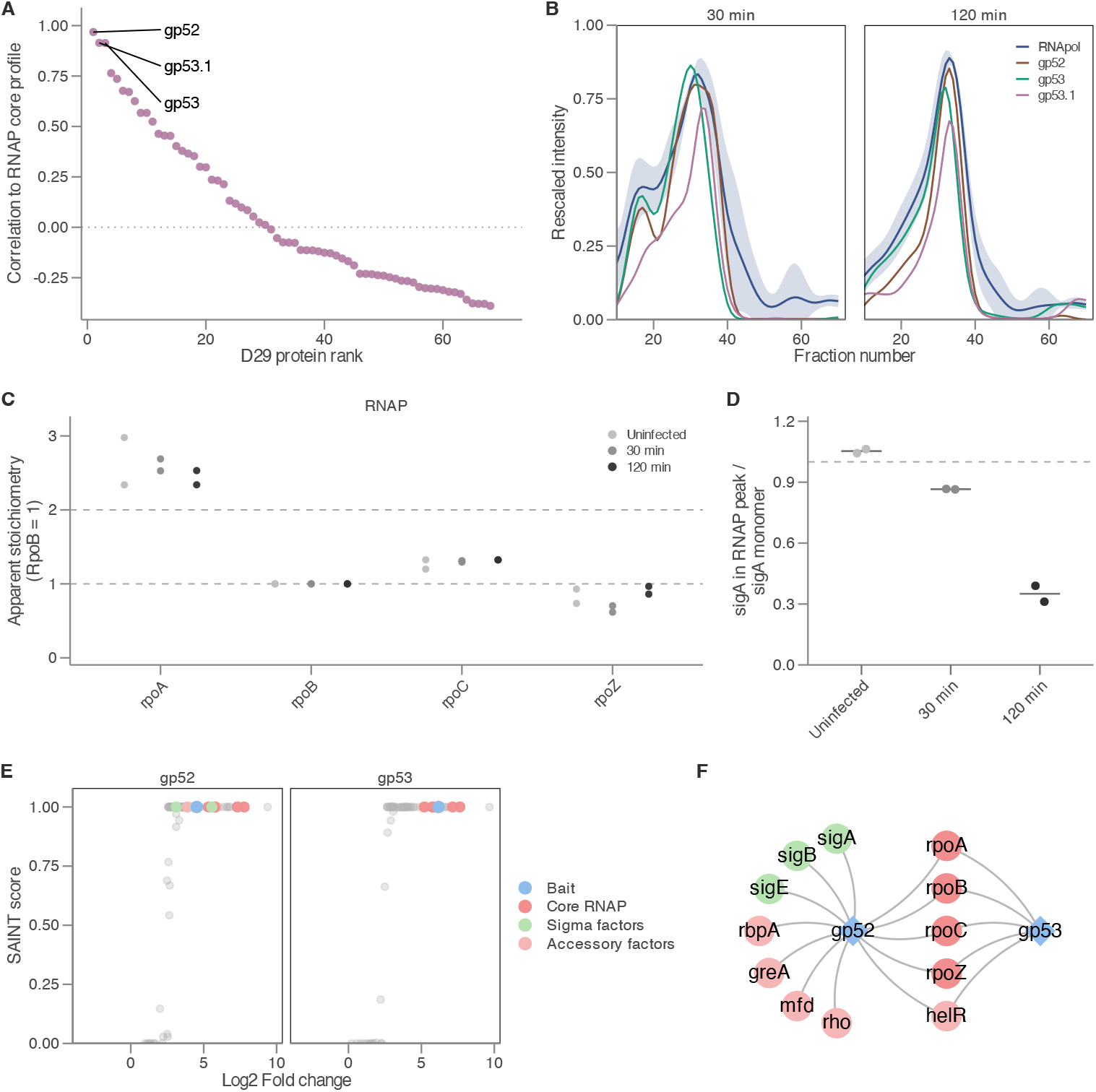
Identification and validation of phage proteins interacting with mycobacterial RNA polymerase. **A.** D29 proteins ranked by co-elution with the RNA polymerase (RNAP) core. Y-axis shows the Pearson correlation between each D29 proteins identified at 30 minutes post infection and the RNAP core profile (the mean of the z-scored profiles of RpoA, RpoB, RpoC and RpoZ) vs its rank (X-axis). **B.** SEC co-elution profiles for RNAP core and its putativep hage interactors. The x-axis shows the fraction number and the y-axis the rescaled intensity. The blue line is the mean profile of the core RNAP subunits (rpoA, rpoB, rpoC and rpoZ), and the shaded band indicates mean *±* SD across subunits and replicates. Coloured lines show the co-eluting phage proteins gp52, gp53 and gp53.1. **C.** Apparent stoichiometry of the core RNAP subunits in the RNAP SEC peak (estimated by apparent molecular weight) in uninfected cells and at 30 and 120 min post-infection. Protein abundances were estimated by iBAQ and normalised to RpoB (= 1). Each dot represents one replicate (*n* = 2). Dashed lines at 1 and 2 mark the expected *α*_2_*ββ^′^ω* stoichiometry. **D.** Ratio of sigA abundance in the RNAP peak to sigA in the monomer peak (fractions 62–70) for each SEC-MS run. The dashed line marks equal amounts. Points are replicates (*n* = 2); bars show the mean. **E.** Scatter plot of SAINT score versus log_2_ fold change for proteins detected in gp52 and gp53 AP-MS in uninfected *M. smegmatis*. Each point represents one protein. Bait proteins, core RNAP subunits, sigma factors (sigA, sigB, sigE) and accessory factors (HelR, RbpA) are highlighted. **F.** Network of medium and high-confidence interactions (SAINT score *>* 0.8) from gp52 and gp53 AP-MS in uninfected *M.smegmatis*. Diamonds represent phage proteins and circles host proteins; colours as in E.

### Evolutionary conservation and diversification of gp53–gp52

We next examined the evolutionary conservation of gp52 and gp53 to gain insight into their potential molecular functions. In phages, genes that are functionally related are often conserved within defined lineages, and comparative analysis has proven informative for mechanistic prediction [27]. We therefore analyzed the phylogenetic distribution and sequence architecture of gp52 and gp53 across all sequenced mycobac-teriophages, supplemented with a targeted profile-HMM search (Pfam PF25684 for gp53, PF17468 for gp52; *E <* 1×10*^−^*^10^) against BFVD [28] and proGenomes [29]. This combined search yielded 810 gp53 homologues, including 789 from Phamerator/SEA-PHAGES mycobacteriophages and 21 from BFVD and proGenomes, which were clustered with MMseqs2 at 80% amino-acid identity into 81 non-redundant representa-tive clusters for phylogenetic analysis (Fig. 3A). Within Cluster A, both gp53 and gp52 were detected in every genome surveyed (*n* = 788, 100% conservation), indicating that the two proteins constitute a conserved feature of this lineage. We additionally identi-fied gp53.1 as a highly conserved but more lineage-restricted component of the locus. Profile-HMM searches identified 369 gp53.1 homologues, all 37–63 aa in length and confined to Cluster A mycobacteriophages, with gp53.1 being nearly universal within the D29-containing A2 subcluster. The broader gp53 search additionally identified a homologue in the non-mycobacteriophage *Rhodococcus* phage Lillie and in 20 bacterial genomes, predominantly Actinobacteria closely related to *Mycobacterium*, together with one more distant hit in *Paraburkholderia bengalensis* (Fig. 3A). HMMER analy-sis against Pfam-A identified the gp53-associated HTH/DNA-binding family PF25684 in all 81 gp53 representatives, irrespective of locus architecture (Fig. 3A), supporting a conserved DNA-binding function. By contrast, gp52 homologues matched PF17468 (DUF) but showed no detectable HTH-associated Pfam domain. Across the full homo-logue sets, the PF25684 helix-turn-helix region covers almost the entire gp53 protein and the PF17468 domain of unknown function almost the entire gp52 protein, and both regions are well conserved along their length (Supplementary Fig. S4). Mea-sured instead as percent amino-acid identity to the D29 proteins, the two differed with gp52 generally showing stronger conservation across homologues than gp53 (Fig. 3A), consistent with distinct evolutionary constraints on the two components.

**Fig. 3.**
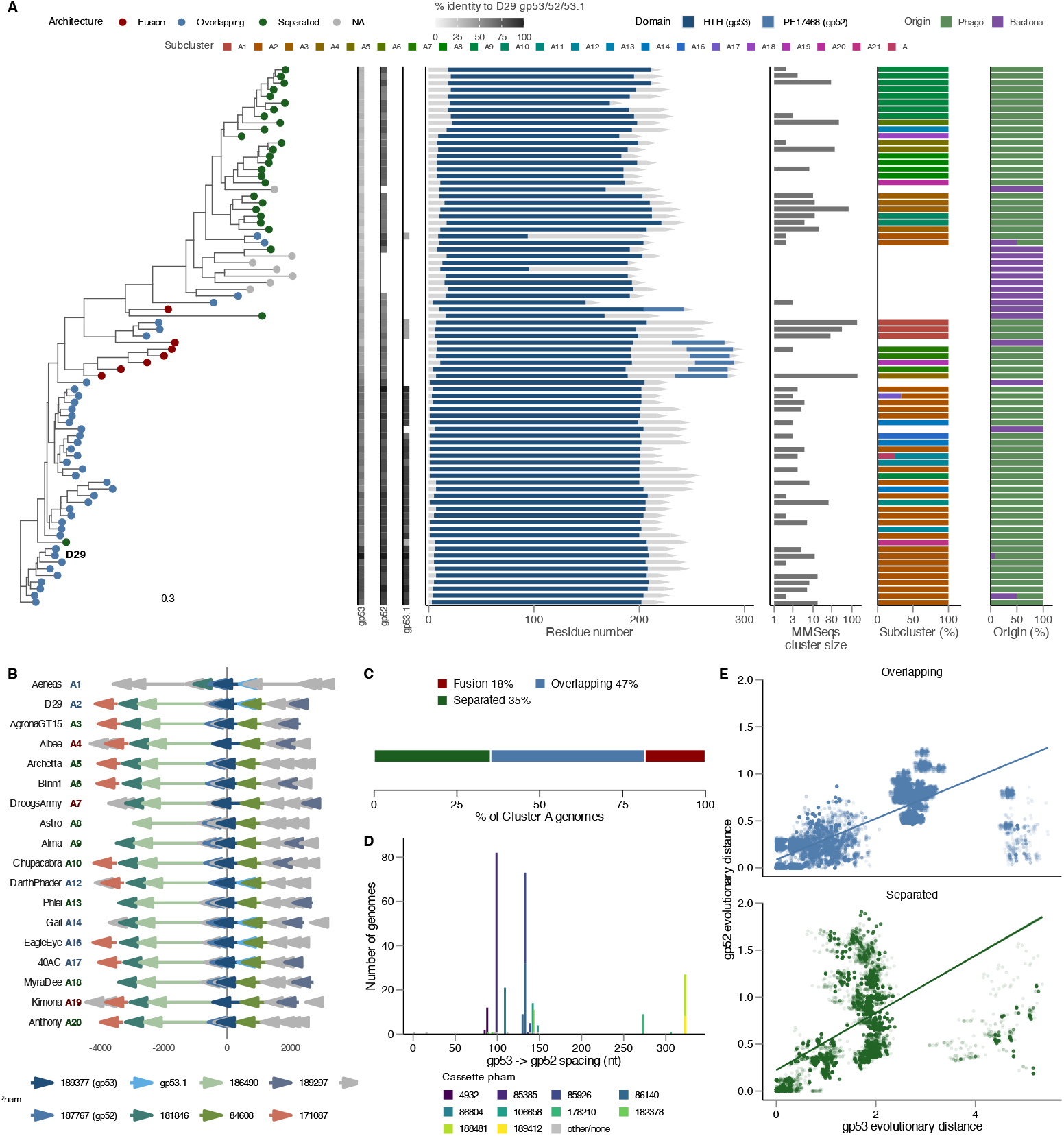
Evolutionary relationships, domain architecture and conservation of gp53 across phage subclusters. **A.** Maximum-likelihood tree of 81 gp53 cluster representatives (MMseqs2, 80% identity, from 810 sequences), D29 labeled, with row-aligned tracks for architecture class, percent identity to D29 gp53/gp52/gp53.1, domain schematic, cluster size (log scale), and subcluster and origin composition (%). **B.** Synteny of gp53 neighbourhoods (*±*5 ORFs) by architecture class. **C.** Architecture class across all 788 Cluster A genomes. **D.** Distribution of the gp53-to-gp52 spacing across the 277 genomes with separated loci, at single-nucleotide resolution. Bars are stacked and coloured by the pham of the first intervening gene; the 10 phams seen in at least five genomes are coloured individually, and singleton phams and genomes with no assignable intervening pham are grey. **E.** Mirrortree: pairwise gp52 versus gp53 patristic distance across 644 genomes (Pearson *r* = 0.61, permutation *p ≈* 2 *×* 10*^−^*^4^).

Within Cluster A, gp53 sequences segregated into clades that closely tracked estab-lished subcluster classifications (Fig. 3A). A similarly subcluster-resolved topology was observed for gp52 (Supplementary Fig. S3), indicating that both proteins largely fol-low the broader evolutionary structure of Cluster A. We next examined the genomic context of the locus across Cluster A phages. Synteny was highly conserved, with gp53 (Pham 189377) and gp52 (Pham 187767) embedded within a stable cassette flanked by phams 181846, 186490, 84608 and 189297, with pham 171087 retained in a sub-set of genomes (Fig. 3B). Where present, gp53.1 was consistently located immediately downstream of gp53, further supporting its association with this genomic module. The gp53/gp52 locus resolved into three recurrent architectures across Cluster A: gp53–gp52 fusion proteins, overlapping gp53 and gp52 ORFs, and loci in which the two genes were separated by one or more ORFs. Fusion accounted for 18% of Cluster A genomes, overlapping architectures for 47% and separated architectures for 35%, with all Cluster A phages having one of these three systems (Fig. 3C). Fusion was strongly lineage-associated, with A4, A7 and A19 encoding the fused form, whereas the remaining represented subclusters predominantly retained distinct but conserved gp53 and gp52 ORFs. Gp53.1 was detected in almost all the overlapping 52/53 archi-tectures (363/367), while not detected in the fused (0/144) or separated (2/277) architecture. Importantly, the separated architecture did not reflect random expan-sion of the gp53–gp52 loci. Instead, across the 277 genomes with separated loci the gp53-to-gp52 spacing was confined to 20 discrete lengths, clustered between 85 and 148 nt and again between 273 and 323 nt (Fig. 3D). The intervening sequence car-ried a varied but bounded set of genes: 15 distinct phams, of which 10 recurred in five or more phages and together accounted for 269 of the 274 genomes with an assignable intervening pham, the remaining five phams being singletons. Gap content was largely fixed for a given spacer length: for 7 of the 10 lengths observed in at least five genomes, ≥90% of genomes carried the same pham or pham combination.

The main exception was the 133-nt spacer, which was split by subcluster between two phams (A6, pham 85926; A9, pham 86140). All 37 genomes with long spacers (273–323 nt; subclusters A5 and A8) encoded one of three Tad1-like phams carrying an Acb2/Tad1 hairpin domain (PF24729). In 33 of these, the Tad1-like ORF was paired with a small 29–36-aa ORF (pham 182378) as a two-gene cassette. By contrast, 237 of the 238 genomes with 85–148-nt spacers encoded a single small ORF (29–60 aa) with no detectable Pfam domain, whose pham varied with subcluster. Thus, variation in gp53–gp52 spacing reflects recurrent lineage-specific cassette architectures rather than stochastic insertion between the two genes. Consistent with this physical linkage, gp53 and gp52 showed significant co-diversification across 644 paired homologues (Mir-rorTree *r* = 0.61, permutation *P* ≈ 2× 10*^−^*^4^) and co-occurred substantially more often than expected by chance (Fisher’s exact *P* = 8.5 × 10*^−^*^255^, odds ratio ≈1,200). Their individual phylogenies nevertheless showed substantial topological discordance (nor-malized Robinson–Foulds distance = 0.92), indicating correlated diversification within a conserved genomic module without identical evolutionary histories. Altogether, these analyses identify gp53 and gp52 as the conserved core of a highly structured Cluster A locus, with the small gp53.1 protein representing a lineage-restricted extension of this module. gp53 retains a universal HTH-associated domain consistent with DNA binding, whereas gp52 belongs to a distinct, more sequence-constrained family with no detectable canonical DNA-binding motif. Gp53 and gp52 remain tightly linked through fusion, overlapping coding sequences or recurrent accessory-gene cassettes, supporting their evolution as a shared functional module while allowing substantial remodeling of the genomic architecture connecting them.

### gp53 binds to Mycobacteria RNAP and displaces SigA

Having established that gp52 and gp53 are encoded within a conserved genomic locus in Cluster A phages and both associate with the host RNA polymerase during infec-tion, we next predicted the structure of RNAP with AlphaFold3 [30] together with gp52 and gp53 to evaluate how these factors might engage RNAP. Motivated by our AP-MS recovery of non-canonical sigma factors, we first modeled gp52 and gp53 in complex with the RNAP holoenzyme (core RNA polymerase composed of *α*_2_*ββ^′^ω* subunits associated with a single *σ* factor, typically *σ^A^* in mycobacteria, forming *α*_2_*ββ^′^ω*–*σ*) to define a structural reference for their mode of interaction, specifically assessing potential occupancy of the sigma-factor binding pocket and the poten-tial rewiring of transcription. Housekeeping sigma factors have three DNA-binding domains: Domain 2 (D2) binds the -10 element to initiate promoter melting, Domain 3 (D3) binds the extended -10 element, and Domain 4 (D4) binds the -35 element[16]. We modeled gp52 and gp53, individually or together, against Mtb RNAP core using AlphaFold3 [30] (Fig. 4A). We then superimposed these structures onto a previously resolved mycobacterial RNAP transcription initiation complex containing promoter DNA (6EDT) [31]. Gp52 was placed near the downstream DNA exit channel (Fig. 4A). As for gp52, its position doesn’t suggest a function directly, but its binding to both *M. smegmatis* sigma factors and RNAP together with extensive conservation, suggest a potential role in rewiring transcription in mycobacteria, which will require follow-up functional assays. On the other hand, gp53 was predicted to bind in the same pocket as SigA domains 2 and 3 (SigA-D2 and SigA-D3), so we termed these domains gp53-D2 and gp53-D3 (Fig. 4B–D). We note that we had to manually adjust the AlphaFold model of gp53-D3 to better align with SigA-D3 (Fig. 4C). These assignments also allow the existing annotation of gene 53 to be evaluated directly. In mycobacteria, SigK (Rv0445c) is an extracytoplasmic function (ECF, group IV) *σ* factor and, like all ECF factors, comprises only regions 2 and 4, held inactive by the membrane-associated anti-*σ* RskA; in B. subtilis, *σ*K is a sporulation-specific factor whose gene is recon-stituted by site-specific excision of the skin element from two split ORFs and whose product requires proteolytic maturation from pro-*σ*K. Neither description fits gp53, which is a single 235-residue protein carrying *σ*2- and *σ*3-like domains, a combination found in primary rather than ECF *σ*factors, and which engages the core enzyme at the *σ*A D2-D3 surface. The *σ*-like character of gp53 is therefore real but class-specific: it mimics the promoter-engaging module of a housekeeping *σ* factor, not the *σ*2-*σ*4 architecture of *σ*K.

**Fig. 4.**
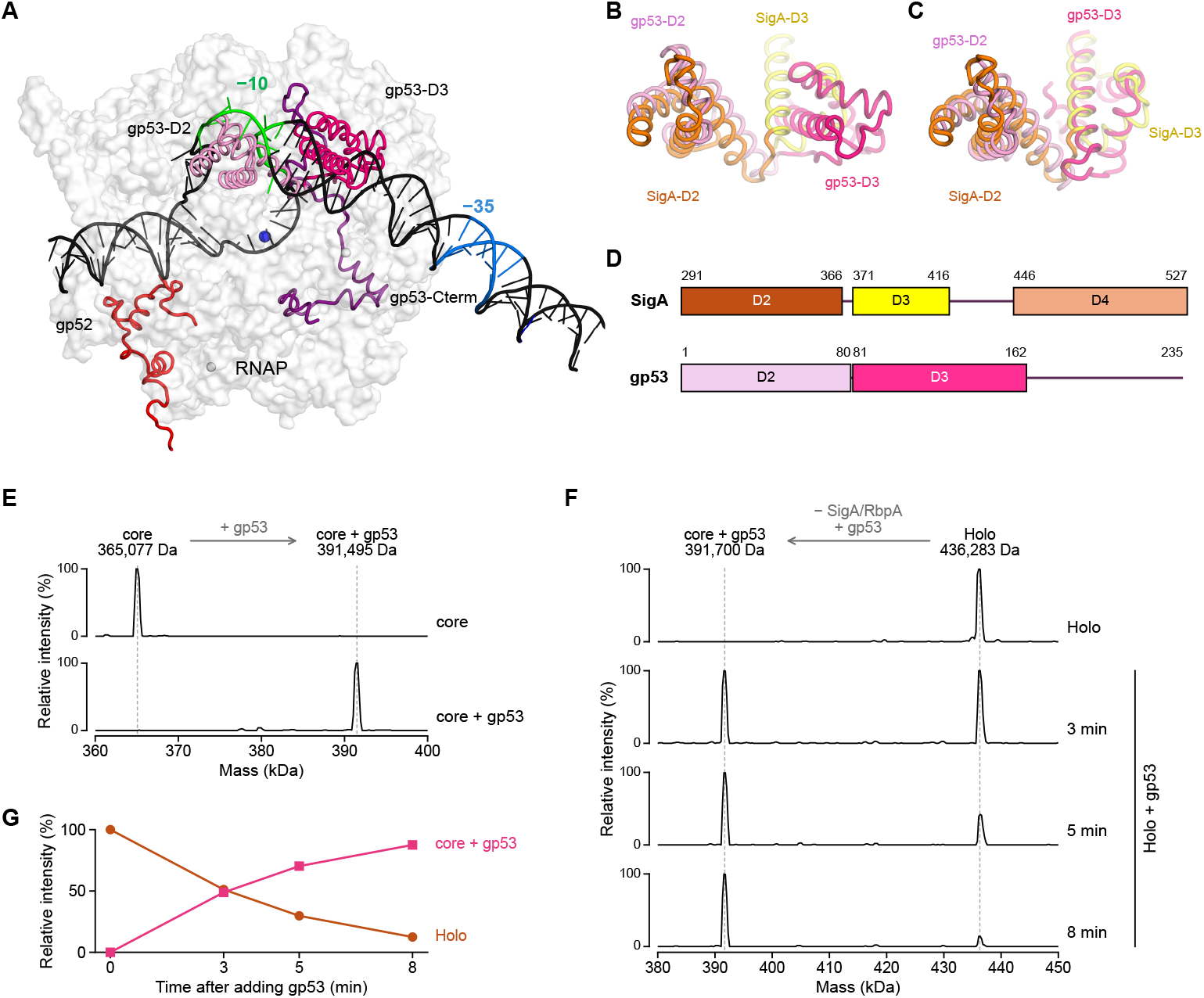
Structural similarities between D29 gp53 and Mtb SigA, and nMS analysis of gp53 binding to Mtb RNAP core and holoenzyme complexes. A. AlphaFold3-predicted models of gp52 (red) and gp53 domains D2 and D3 (pink and magenta, respectively) positioned on the core RNA polymerase open promoter complex (PDB ID: 6EDT). The *−*10 element on the non-template strand is green and the *−*35 element is blue; the blue sphere represents the active-site Mg^2+^. **B.** AlphaFold3-folded gp53 domains D2–D3 on RNAP superimposed on SigA domains D2–D3 on RNAP (6EDT). RNAP is not shown and the view is the same as in A but magnified. **C.** Manual alignment of gp53 domains D2–D3 to SigA domains D2–D3. gp53-D2 aligns well with SigA-D2, but gp53-D3 was made a separate rigid body and manually aligned to SigA-D3. **D.** Domain architecture of Mtb SigA and D29 gp53 drawn on a shared residue axis, aligned at the start of D2. Colours are as in A–C. **E.** Deconvolved nMS spectra of the Mtb RNAP core alone and incubated with gp53 at a 1:1 molar ratio. Dashed lines mark the core (365,077 Da) and core + gp53 (391,495 Da) species. Each spectrum is scaled to its most intense peak. **F.** nMS monitoring of the exchange between host SigA/RbpA and phage gp53 following addition of gp53 to the reconstituted Mtb RNAP holoenzyme at a 1:1 molar ratio. Representative deconvolved mass spectra are shown for the holoenzyme alone and at the indicated times after mixing. Dashed lines mark the core + gp53 (391,700 Da) and holoenzyme (436,283 Da) species. **G.** Relative intensity of the holoenzyme and core + gp53 species over the time course in F, each expressed as a percentage of their combined deconvolved intensity

Based on the predictions, we expect gp53 to interact with the -10 element of the pro-moter. The D29 genome contains several sites known as stoperators which resemble the canonical -35 elements[32, 33], however, these are not paired with the canoni-cal -10 elements. This suggests that D29 might have unique -10 elements which are specifically recognized by gp53. We note that although AlphaFold did not predict the C-terminus of gp53 to bind the same region as SigA-D4, it may still bind the same region of RNAP and recognize the -35 element. However, both functional and struc-tural studies are needed to validate that gp53 is a functional analogue of sigmas. To confirm our SEC-MS and AP-MS data we tested if gp53 binds to mycobacterial RNAP, using purified D29-gp53, *M. tuberculosis* (Mtb) core RNAP, and Mtb SigA/RbpA. We used native mass spectrometry (nMS) to establish the binding stoichiometry and con-sequences of gp53 interaction with Mtb RNAP core and holoenzyme complexes. nMS resolves noncovalent protein assemblies by mass, enabling distinct RNAP complexes to be distinguished simultaneously within a single spectrum. Analysis of the Mtb RNAP core revealed a 365.1-kDa complex consistent with the expected *α*_2_*ββ^′^ω* subunit stoi-chiometry. Incubation with gp53 at a 1:1 molar ratio quantitatively shifted the RNAP population to a 391.5-kDa complex, corresponding to the association of a single 26.4-kDa gp53 molecule with the RNAP core (Fig. 4E). We next reconstituted the Mtb holoenzyme by incubating the RNAP core complex with SigA/RbpA to test poten-tial competition or displacement of *σ*A by gp53. nMS confirmed the formation of the 436.3-kDa holoenzyme with correct subunit stoichiometry (*α*_2_*ββ^′^ωσ*ARbpA). We then continuously monitored the interaction of gp53 with the preassembled holoenzyme in a time-course experiment by nMS. The SigA/RbpA-bound holoenzyme progressively decreased with a concomitant increase in the gp53-bound RNAP core complex, culmi-nating in complete conversion to the gp53-bound species. These measurements directly captured the exchange of SigA/RbpA for gp53 on RNAP (Fig. 4F,G). Together, the nMS results establish that gp53 binds RNAP with a 1:1 stoichiometry and displaces and replaces the host SigA/RbpA from the RNAP, providing direct biochemical evi-dence that it functions as anti-host sigma, likely by competing for the same binding site as SigA on RNAP.

## Discussion

Bacteriophages encode many conserved proteins whose roles in host takeover remain poorly understood. Transcription is a major control point in this process, particularly for mycobacteriophages, which depend on host RNA polymerase (RNAP) for phage gene expression. Productive infection therefore requires not only access to RNAP, but mechanisms that redirect its activity toward the viral program. Here, unbiased inter-action proteomics identified three D29 proteins, gp52, gp53 and gp53.1, that associate with host RNAP during infection, with these interactions independently recovered by affinity purification. Native mass spectrometry further showed that gp53 alone can displace the housekeeping factor *σ^A^* from purified *M. tuberculosis* RNAP. Together with structural modelling placing gp53 at the sigma-binding surface, these data sup-port a model in which gp53 directly competes with host sigma factors for RNAP. This mechanism is distinct from previously characterized phage strategies for altering sigma-dependent transcription. T4 achieves transcriptional reprogramming through a staged system in which AsiA and MotA first remodel host *σ*^70^ for middle transcription, followed by replacement of *σ*^70^ by the phage-encoded late sigma factor gp55. Notably, gp55 alone is a poor competitor for RNAP and is readily displaced by *σ*^70^. Other phages instead use dedicated inhibitors: Xp10 P7 displaces host sigma during pro-moter engagement and blocks productive transcription initiation, whereas phiEco32 gp79 inhibits *σ*^70^-dependent transcription while promoting transcription by a separate phage-encoded sigma factor, gp36. In contrast, gp53 combines sigma-like architecture with the ability to directly displace *σ^A^* from preassembled mycobacterial RNAP in the absence of an additional phage factor. Despite this activity, gp53 expression does not compromise cell viability or cause global transcriptional shutdown, arguing against a primary role as a general transcriptional inhibitor. Its compact Domain 2- and Domain 3-like architecture instead suggests that gp53 represents a minimal sigma-factor mimic capable of directly competing for the sigma-binding interface. Whether gp53 also con-fers promoter specificity and therefore functions as a bona fide phage sigma factor remains to be established. The evolutionary organization of the locus suggests that this mechanism extends beyond D29. gp53 and gp52 are conserved across Cluster A mycobacteriophages and occur in several recurrent genomic arrangements, including fused proteins, overlapping genes and separated loci containing lineage-specific acces-sory genes. Despite this variation, gp53 retains a conserved HTH-associated domain and gp52 remains tightly linked to it, consistent with an evolutionarily conserved tran-scriptional takeover module that has diversified across phage lineages. More broadly, this study shows how interaction proteomics can resolve functional assignments that sequence analysis alone leaves unsettled. Gene 53 is a case in point: a *σ*-factor label has been carried in mycobacteriophage genome records for decades without experimental support and without a proposed mechanism, while the same sequence carries no *σ*70-family domain assignment. Our data replace that label with a defined and testable mechanism, direct competition with *σ*A for the *σ*-binding surface of RNAP. SEC-MS recovered gp53 and gp52 through their association with endogenous RNAP during infection while preserving major host macromolecular complexes and near-complete coverage of the D29 proteome. These data provide a systems-level view of host machin-ery engagement during infection and identify additional host-phage interactions for mechanistic study. Together, our findings define a conserved mycobacteriophage mod-ule that targets the sigma-binding surface of RNAP and suggest that remodeling of the host polymerase may represent a broader strategy for transcriptional takeover in Cluster A phages.

## Methods

### Growth conditions

Experiments with *M. smegmatis* mc^2^155 (RefSeq Assembly: GCF 000015005.1) cul-tures were grown in Middlebrook 7H9 media (BD-Difco) supplemented with glycerol and Tween 20. Electroporation transformation experiments in *E. coli* used NEB 10-Beta cells (NEB) and recovery with Stable Media, and plasmids were selected under 50 µg/ml of kanamycin sulfate (Sigma). Subsequently, *E. coli* were grown in Luria Bertani broth [34].

### Plaque assay

Plaque assays were performed at 37 *^◦^*C using solid Middlebrook 7H9 media (BD-Difco) MB7H9 agar plates. A total of 150 µL of overnight *Mycobacterium smegmatis* culture was mixed with 3 mL of top agar (0.35% MB7H9 agar) and poured onto a solidified bottom agar plate (20 mL MB7H9 agar). Serially diluted phage lysates were spotted (5 µL) onto the top agar after solidification. Plates were incubated at 37 *^◦^*C, and plaques were enumerated after 24 h.

### Bacterial culture and infection

*M. smegmatis* strain mc^2^155 was grown overnight in 5 mL MB7H9 medium at 37 *^◦^*C with aeration (175 rpm). Cells were diluted 1:100 from a saturated overnight culture into 100 mL of fresh MB7H9 and grown to mid-logarithmic phase (OD_600_ = 0.5–0.6; ≈ 3 × 10^8^ CFU/mL). Cultures were infected with mycobacteriophage D29 at a multiplicity of infection (MOI) of ∼10 and incubated for 10 min at 37 *^◦^*C to allow complete virion adsorption. Following adsorption, cultures were further incubated for 20 min or 110 min, corresponding to total infection times of 30 min and 120 min, respectively.

Cells were harvested by centrifugation at 6,000 × *g* for 5 min at 0 *^◦^*C. The supernatant was discarded, and pellets were washed twice with 5 mL of ice-cold MB7H9. The final cell pellets were combined for downstream size-exclusion chromatography (SEC) sample preparation.

### Mtb RNAP complex purification

Mtb RNAP was purified following a previously established protocol [31]. Briefly, pMP61 plasmid containing the genes for the Mtb core subunits under control of the T7p was transformed into *E. coli*Rosetta2 DE3 cells. Cells were grown at 37 *^◦^*C to an optical density at 600 nm (OD_600_) of ∼0.3, then grown to an OD_600_ of ∼0.6 at 25 *^◦^*C, and RNAP expression was induced using 0.1 mM isopropyl *β*-D-1-thiogalactopyranoside (IPTG; Gold Biotechnology, 1248) overnight at 25 *^◦^*C. Cells were collected by centrifugation, resuspended in lysis buffer (50 mM Tris-HCl, 5% glycerol pH 8.0, 1 mM EDTA, 5 mM dithiothreitol (DTT), 1 mM phenylmethyl sul-fonyl fluoride, 1 mM protease inhibitor cocktail), and lysed by three passages through microfluidizer (Microfluidics). The lysate was centrifuged twice (27,000 × *g*, 15 min, 4 *^◦^*C). Polymin P (PEI, Thermo Fisher, AC178572500) was added to the supernatant to a final concentration of 0.6% (w/v) and stirred for 15 minutes at 4 *^◦^*C. The solution was then centrifuged (11,000 × *g*, 15 min, 4 *^◦^*C) and the pellet was washed three times in TGED wash buffer (10 mM Tris-HCl pH 8.02, 5% v/v glycerol, 0.1 mM EDTA, 5 mM DTT, 300 mM NaCl) using a douncer. RNAP was eluted from the pellet using elution buffer (10 mM Tris-HCl pH 8.02, 5% v/v glycerol, 0.1 mM EDTA, 5 mM DTT, 1 M NaCl) in three consecutive elutions. RNAP was precipitated overnight at 4 *^◦^*C using 3.5 g l*^−^*^1^ pulverized (NH_4_)_2_SO_4_. After centrifugation (11,000 × *g*, 40 min, 4 *^◦^*C), the precipitate was dissolved in Nickel buffer A (20 mM Tris pH 8.0, 5% glycerol, 1 M NaCl, 5 mM *β*-mercaptoethanol, 10 mM imidazole) and loaded onto two HisTrap FF 5 ml columns (GE Healthcare Life Sciences). RNAP was eluted from the column with Nickel buffer B (20 mM Tris pH 8.0, 5% glycerol, 1 M NaCl, 5 mM *β*-mercaptoethanol, 250 mM imidazole). Lastly, eluted RNAP was run over a gel filtration column (HiLoad Superdex 26/600 200 pg) in gel filtration buffer (10 mM Tris pH 8.0, 5% glycerol, 0.1 mM EDTA, 500 mM NaCl, 5 mM DTT). Clean fractions were flash-frozen and stored at −80 *^◦^*C.

### Mtb SigA-RbpA complex purification

Mtb SigA-RbpA was purified as follows: A plasmid overexpressing 10His-SigA-RbpA in a single operon via the T7p was transformed into *E. coli* BL21(DE3) cells. Protein production was induced with 0.5 mM IPTG overnight at 18 *^◦^*C. Cells were resuspended in lysis buffer (20 mM Tris-HCl pH 8.0, 5 mM imidazole, 500 mM NaCl, 5% glycerol, 5 mM BME, 0.5 mM PMSF, 1X protease inhibitor) and lysed by the microfluidizer. Lysed cells were clarified by centrifugation and loaded onto two HisTrap FF 5 ml columns (GE Healthcare Life Sciences). The column was washed with Nickel A buffer (20 mM Tris-HCl pH 8.0, 5 mM imidazole, 500 mM NaCl, 5% glycerol, 5 mM BME) and eluted with Nickel B buffer (20 mM Tris-HCl pH 8.0, 500 mM imidazole, 500 mM NaCl, 5% glycerol, 5 mM BME). Fractions containing the protein were combined and treated with precision X protease overnight at 4 *^◦^*C while dialyzing in Nickel A buffer.

Cleaved protein was purified by reverse Nickel column using the same buffers. Fractions containing the protein were combined and loaded onto a Superdex 200 HiLoad (Cytiva) column and eluted using gel filtration buffer (20 mM Tris-HCl pH 8.0, 500 mM NaCl, 5% glycerol, 0.1 mM EDTA, 1 mM DTT). Clean fractions were flash-frozen and stored at −80 *^◦^*C.

### D29 gp53 purification

The D29-gp53 gene block was synthesized (Twist Bioscience) and cloned into a pET-10His-SUMO plasmid by Gibson assembly. The expression plasmid was transformed into Rosetta 2 (DE3) competent cells and grown to an OD_600_ of 0.6 at 37 *^◦^*C with shaking at 220rpm. Protein expression was induced with 0.5mM IPTG for 3 hours at 30 *^◦^*C. The gp53 protein was then purified as with SigA-RbpA, except cells were lysed by sonication (Branson), protein was treated with UlP1 instead of Precision X protease after the first Nickel column, and a Superdex 75 (Cytiva) increase was used for size-exclusion chromatography.

### Generation of ectopically tagged gp53, gp52

gp52 oligonucleotides were synthesized as gBlocks by IDT. gBlocks included a syn-thetic RBS generated by the RBS calculator (Denovodna.com) [35] designed for maximum translational efficacy in *Mycobacterium smegmatis* under the acetamide-inducible promoter in pUS116. gBlocks also included both C- and N-terminal Strep-II tags, respectively. Briefly, assembly of Strep-II-tagged gp52 and gp53 was performed using NEB’s HiFi NEBuilder Mastermix into shuttle vector pUS116 (Addgene 84578) [36] by PCR amplifying the vector with 35 bp overhangs for the gBlock sequence and incubated at 50 *^◦^*C for 1 h. Assembled plasmids were diluted 1:1 in ddH_2_O and 1 µL was electroporated into NEB 10-Beta cells following the manufacturer’s proto-col and selected under 50 µg/ml kanamycin sulfate. Plasmids were purified using the Monarch Plasmid Miniprep kit (NEB). Subsequently, whole-plasmid sequencing (Plas-midsaurus) was performed and validated plasmids were transformed into *M. smegmatis* following standard protocols [37].

### Affinity purification of tagged phage proteins

*M. smegmatis* transformed with pUS116 synRBS gp52 ctermSII, pUS116 synRBS gp52 ntermSII and pUS116 empty vector were grown from single colonies overnight in 7H9 media (36 h) under kanamycin selection. 50 mL (for each replicate, plasmid and empty vector — 450 mL total) of 7H9 media supplemented with kanamycin was then inoculated with actively growing cultures 1:100 and grown for an additional 18 h. Exponentially growing cells (OD_600_ 0.8) were induced with acetamide. Acetamide was prepared at 30% w/v in water and filter sterilized (0.22 µm) before adding to a final concentration of 0.3% w/v for induction. After 18 h of induction, cells were centrifuged for 30 min at 12,000 × *g* at 4 *^◦^*C and resuspended in 10 mL of TNN lysis buffer (1% NP-40, 100 mM Tris-HCl, 150 mM NaCl, 1 mM EDTA and Roche cOmplete Ultra Protease Inhibitor cocktail, 600 nM avidin from IBA BioLock, pH 8.0). Cells were loaded in 2 mL tubes containing soda glass beads. Cells were lysed via a FastPrep24 homogenizer (MP Biomedicals) at 10 m/s for 4 rounds of 45 s with 2 min on ice in between each round. Lysates were clarified by centrifugation (maximum speed, 17,000 × *g* at 4 *^◦^*C) for 30 min.

For purification, 500 µL 50% Strep-Tactin XT 4Flow high-capacity (IBA) slurry was applied to Bio-Rad empty columns and allowed to equilibrate at 4 *^◦^*C for 30 min. Columns were washed in 10 consecutive column volumes (250 µL/wash) of the lysis buffer without allowing the column to run dry. After washing the column, clarified and diluted lysate was applied to the column and allowed to pass entirely through the column. The column was washed in lysis buffer in 10 consecutive washes without allowing the column to run entirely dry. Elution was performed with 1× Buffer BXT (IBA) supplemented with protease inhibitor cocktail in 3 column volumes. All purifi-cation steps were performed at 4 *^◦^*C in a cold room. TCA was added to 25% final concentration to precipitate the eluted proteins and stored at −20 *^◦^*C until proteomics sample processing.

### Sample preparation for SEC–DIA–MS and bulk proteomics

*M. smegmatis* pellets infected with D29 at 0, 30 and 120 min were resuspended in 500 µL SEC running buffer (50 mM HEPES, 150 mM NaCl, pH 7.4) supplemented with protease inhibitors (Roche). The resulting supernatant was mechanically lysed using steel beads and a bead beater in HN buffer. Samples were ultracentrifuged for 30 min at 4 *^◦^*C to remove debris. Lysates were concentrated to 100 µL using a 30 kDa molec-ular weight cutoff filter to enrich for high-molecular-weight assemblies and deplete monomeric proteins. The concentrated material was centrifuged at 10,000 × *g* for 10 min at 4 *^◦^*C, and the clarified supernatant was collected for chromatographic separation. 100 µg were reserved for bulk proteomics.

### Size-exclusion chromatography

Approximately 1,000 µg of total protein (as estimated by BCA assay) was injected on an Agilent Infinity 1260 HPLC system operating at 0.5 mL/min in SEC running buffer. Chromatographic separation was performed using a Yarra Phenomenex SRT-C1000 column maintained at 4 *^◦^*C. 70 fractions (100 µL each) were collected between 8–22 min. The column was washed with two column volumes (10 mL) of SEC buffer between runs. Molecular weight calibration was performed using the Phenomenex AL0-3042 protein standard mixture (100 µL protein mix + 6 µL uridine).

### SEC–MS proteomics sample preparation

SEC fractions were processed using a 96-well filter-aided sample preparation (FASP) protocol as previously described [24]. Filters were washed twice with 100 µL ddH_2_O.

SEC buffer was removed by centrifugation (1,800 × *g*, 1 h, RT), and proteins were resuspended in 50 µL of TUA buffer (5 mM TCEP, 8 M urea, 20 mM ammonium bicar-bonate) and incubated at 37 *^◦^*C for 30 min with agitation (400 rpm). Cysteines were alkylated by adding 20 µL CAA buffer (35 mM chloroacetamide, 20 mM ammonium bicarbonate) and incubating for 1 h at 25 *^◦^*C in the dark.

Following reduction and alkylation, TCEP and CAA were removed by centrifugation (1,800 × *g*, 30 min), and filters were washed three times with 100 µL of 20 mM ammo-nium bicarbonate. Proteins were digested in 50 µL of 20 mM ammonium bicarbonate containing 1 µg of trypsin per fraction and incubated overnight. Peptides were col-lected by centrifugation into a 96-well receiver plate (1,800 × *g*, 30 min) and washed once with 100 µL ddH_2_O. Filters were then dried by centrifugation (1,800 × *g*, 60 min).

Combined peptides from each fraction were vacuum-dried and stored at −80 *^◦^*C until LC–MS acquisition.

### AP-MS sample preparation

Dried proteins were resuspended in 100 µL of 8 M urea, 100 mM ammonium bicar-bonate (ABC, pH 8.1). Samples were reduced with 5 mM TCEP for 30 min at room temperature, and cysteines were alkylated with 10 mM chloroacetamide (CAA) for 30 min in the dark. Urea concentration was diluted to 1 M with 100 mM ABC, and proteins were digested with 2 µg of trypsin for 14 h at 37 *^◦^*C (600 rpm). Reactions were quenched by adding 10% formic acid (FA), and peptides were desalted using C18 spin columns (Nest Group). Columns were activated with 1 CV ACN, equilibrated with 2 CV 0.1% FA, and loaded twice with peptide solution. After washing with 3 CV 0.1% FA, peptides were eluted with 0.5 CV of 50% ACN/0.1% FA (repeated twice). Samples were dried under vacuum and stored at −80 *^◦^*C until LC–MS acquisition.

### AP-MS data acquisition

AP-MS samples were acquired on a timsTOF Pro2 connected to a ProteoElute UHPLC (Bruker). Peptides were separated at 300 nl/min on a PepSep column (25 cm × 75 µm × 1.5 µm) heated at 50 *^◦^*C using a linear gradient from 5% buffer B (0.1% FA in 90% ACN) to 33% B in 50 min; buffer B was then increased to 45% in 5 min and the column was washed at 100% B for 10 min. The samples were acquired in DDA-PASEF with the following settings: IM range (0.85–1.3 1/K_0_) and MS1 range 100–1700. 4 PASEF ramps were used (cycle time 0.5 s) and ions with a minimal intensity of 20,000 and max charge of 5 were fragmented. Fragmented precursors were dynamically excluded for 0.4 min. Capillary voltage was set at 1600 V and 180 *^◦^*C.

### Native MS analysis

The Mtb holoenzyme complex was reconstituted by mixing RNAP core complex and *σ*A-RbpA at 1:2 molar ratio and incubated at 37 *^◦^*C for 15 min. The RNAP core complex, holoenzyme complex and gp53 samples were buffer-exchanged twice into 500 mM ammonium acetate pH 7.5, 0.01% Tween-20 using Zeba microspin desalting columns with a 7-kDa molecular weight cutoff (Thermo Scientific). A 3-µL aliquot of the buffer-exchanged sample was loaded into a gold-coated quartz capillary tip and then electrosprayed into an Exactive Plus with extended mass range (EMR) instru-ment (Thermo Fisher Scientific) with a modified static direct infusion nanospray source [38]. For the core + gp53 sample, 2 µM RNAP core was mixed with 2 µM gp53 and incubated at RT for 5 min prior to buffer exchange and nMS characterization. For the continuous monitoring time course experiment with nMS readout, a 10-µL mixture containing 1 µM each of the buffer-exchanged holoenzyme and gp53 was prepared and immediately loaded into the conductive tip for subsequent nMS analysis. The nMS parameters used were spray voltage, 1.21 kV; capillary temperature, 150 *^◦^*C; in-source dissociation, 20 V; S-lens RF level, 200; resolving power, 8,750 at *m/z* of 200; AGC target, 1 × 10^6^; maximum injection time, 200 ms; number of microscans, 5; total number of scans, at least 50. Additional nMS parameters were injection flatapole, 7 V; interflatapole, 4 V; bent flatapole, 4 V; high energy collision dissociation (HCD), 200 V; ultrahigh vacuum pressure, 7.2 × 10*^−^*^10^ mbar. Mass calibration in positive extended mass range mode was performed using cesium iodide. For data processing, the collected nMS spectra were visualized using Thermo Xcalibur Qual Browser (v. 4.8.67.0). Spectral deconvolution was performed using UniDec v. 4.2.0 [39, 40] with the following general parameters: smooth charge state distribution, enabled; peak shape function, Gaussian; Degree of Softmax distribution (beta parameter): 5–10. For the time-course experiment, MS scans were averaged over 1-min intervals (∼50 scans) before deconvo-lution. Manual sample loading, source setup and initiation of electrospray ionization resulted in an approximately 2-min delay between mixing the holoenzyme with gp53 and the start of MS data acquisition. Accordingly, the earliest analyzed time point was 3 min after mixing, comprising the 2-min acquisition delay followed by the first 1-min interval of averaged MS scans. The expected masses for the component proteins of the Mtb RNAP holoenzyme complex include *α*: 37,706.4 Da, fused *β* and *β*’: 277,907.0 Da (N-terminal methionine removed, includes two Zn^2+^ ions with each ion coordinated to four cysteines and one Mg^2+^ ion), *ω*: 11,768.1 Da (N-terminal methionine removed), *σ*A : 58,091.5 Da and RbpA: 12,840.5 Da. The expected mass for gp53 is 26,426.1 Da. The observed mass deviation from the nMS measurements of the observed complexes (calculated as the percentage difference between the measured and expected masses relative to the expected mass) ranged from 0.001–0.06%.

### Proteomics data analysis

All DIA samples were searched with DIA-NN (v1.8) [41] using a library-free approach. Files were searched using a combined *Mycolicibacterium smegmatis* mc^2^155 proteome (6731 entries) supplemented with the D29 genome entry downloaded from NCBI and annotated in Phold [42] using --foldseek_gpu and --ultra_sensitive flags (full parameters in log). DIA-NN MS1 and MS2 tolerances were set to 5 and 10 ppm, respectively. Library generation parameters were as follows: charge state precursors (2–5), 1 missed cleav-age, MS1 mass range matching the DIA acquisition scheme. Protein grouping was performed based on the library protein IDs and cross-run normalization was dis-abled. Following the search, the global report file was filtered to ≤1% protein group Q-values (‘Lib.PG.Q.Value’) to derive the final protein number. AP-MS raw files were searched with FragPipe[43] (v23.1; MSFragger v4.3, MSBooster v1.3.17, Perco-lator v3.7.1, Philosopher v5.1.2, IonQuant v1.11.11) using the LFQ-MBR workflow. Default parameters were used, except that match-between-runs was disabled to avoid transferring ids without MS2-based identification. The FASTA database contained the *Mycolicibacterium smegmatis* mc^2^155 proteome (6731 entries), the Strep-tagged bait constructs and the D29 phage proteome (77), plus common contaminants and reversed decoy sequences. Enzyme specificity was strict trypsin (cleavage after K/R, including before P). Up to two missed cleavages were allowed, with peptide lengths of 7-50 residues. Precursor and fragment mass tolerances were both ±20 ppm, with mass calibration and parameter optimization enabled and isotope errors of 0/1/2 allowed. Carbamidomethylation of cysteine (+57.0215 Da) was a fixed modification. Methionine oxidation (+15.9949 Da) and protein N-terminal acetylation (+42.0106 Da) were variable modifications, with at most three variable modifications per pep-tide, and initiator methionine clipping was enabled. MSBooster rescored the PSMs using deep-learning-predicted retention times and fragment spectra, and Percolator validated them. Protein inference used ProteinProphet. Philosopher filtered results to 1% FDR at the PSM, peptide and protein levels (sequential filtering, picked pro-tein FDR). Label-free quantification was done with IonQuant using unique and razor peptides, with MaxLFQ enabled (minimum one ion), 10 ppm m/z tolerance, 0.4 min retention time tolerance and normalization across runs.

### Protein complex prediction from SEC data

Prediction of protein complexes was performed using PCprophet (https://github.com/anfoss/PCprophet/tree/dev-branch, revision #8737964) [21]. Briefly, as reference database *M. smegmatis* STRING (filtered at *>*500 inter-action score) [44] was used, FDR was set to 0.5 and protein complex hypotheses were collapsed to the maximum probability per subcomplex (-co ’PROB’). All other parameters were used as default.

### Bulk proteomics data analysis

Protein quantification and normalization were performed using MSstats (v4.6.1; https://msstats.org) [45]. The peptide-level matrix generated from DIA-NN was imported into R and reformatted to the MSstats long format, containing protein, pep-tide sequence, precursor charge, run, and intensity information. Data were processed using the dataProcess function with normalization set to equalizeMedians, feature summarization to all, and removal of proteins with more than 50% missing values within a condition (remove50missing = TRUE). Missing values were treated as not censored (censoredInt = "NA") and imputation was disabled (MBimpute = FALSE). Protein intensities were log_2_-transformed and summarized to protein-level abun-dances using the robust median polish algorithm implemented in MSstats. All other parameters were used as default.

### AP-MS data analysis

A Python implementation of SAINT (https://github.com/FossatiLab/saintPy, rev #eba56ac) [46, 47] was used to identify significant interactions. Briefly, spectral counts from the combined_protein.tsv matrix from FragPipe were converted to long format and utilized as is without further preprocessing. SAINT was run with default parameters (f = 0, L = 100, R = 100). Interactors were considered significant with a SAINT score *>*0.8 as we previously reported [23, 26].

### Phylogenetic analysis of gp53 and gp52 protein families

Protein-coding gene sets for all sequenced actinobacteriophages (2,474 genomes; 275,501 proteins), together with pham (phamily) and SEA-PHAGES cluster/sub-cluster assignments, were obtained from Phamerator [48] (retrieved 2024). For each target gene the analysis set was the union of (i) all Phamerator-curated pham mem-bers (phams 189365 and 189377 for gp53, pham 187767 for gp52, excluding the gp52–gp53 fusion pham 177529) and (ii) additional homologs from an uncapped MMseqs2 [49] easy-search of the D29 query protein against the full phage proteome (--search-type 1 --cov-mode 2 -c 0 -s 7.5 --max-seqs 20000), retained at ≥25% identity and *E* ≤ 1 × 10*^−^*^3^. This union gave one gp53 and one gp52 protein in each of the 788 Cluster A genomes. The gp53 set was further supplemented with 21 homologues recovered by an HMMER v3.4 [50] search of BFVD [28] and proGenomes [29] for the PF25684 (gp53) and PF17468 (gp52) Pfam [51] families (*E* ≤ 1 × 10*^−^*^10^). Including the D29 query, the resulting 810 gp53 and 789 gp52 sequences were reduced to representatives at 80% identity with MMseqs2 easy-cluster (--min-seq-id 0.8 -c 0.8 --cov-mode 0 -s 7.5), giving 81 gp53 and 30 gp52 tips.

Representative sequences were aligned with MAFFT v7.490 (--auto) [52] and trimmed with trimAl v1.4.1 (-automated1) [53]. For the family conservation tracks (Supplementary Fig. S4) the full 810 gp53 and 789 gp52 sequence sets were aligned with MAFFT (--auto) without trimming. For each alignment column, occupancy was taken as the fraction of sequences carrying a residue rather than a gap, and conserva-tion as 1 − *H/* log_2_ 20, where *H* is the Shannon entropy over the non-gap residues of that column; columns below 50% occupancy were masked. Pfam domain extents were obtained by hmmscan of the D29 protein against the PF25684 (gp53) and PF17468 (gp52) models and mapped onto alignment coordinates through the D29 row, giv-ing residues 1–225 of 235 for gp53 (domain i-Evalue 2.4 × 10*^−^*^82^) and 1–59 of 61 for gp52 (1.4 × 10*^−^*^45^). Maximum-likelihood trees were inferred with IQ-TREE v3.0.1 [54] under ModelFinder [55] restricted to LG/JTT/VT/WAG (best-fit LG+F+G4 for gp53, LG+G4 for gp52), with 1,000 ultrafast-bootstrap [56] and 1,000 SH-aLRT [57] replicates and --keep-ident. Trees were processed in R v4.3.1 with ape [58] and ggtree [59] to relabel tips, deduplicate genomes, and join Phamerator gene, cluster and architecture metadata; each representative was annotated with the size of its 80%-identity cluster and the SEA-PHAGES subcluster composition of all its members. Per-representative percent identity to D29 gp53 and gp52 was computed by MMseqs2 easy-search (--alignment-mode 3), with a Biostrings global-alignment fallback where no MMseqs2 seed hit was found. HTH (gp53) and gp52 domain spans were assigned by hmmscan against Pfam-A (HMMER v3.4), keeping hits with independent E-value *<* 0.01 to PF25684 (Mycobacteriophage Gp53) and PF17468 (GP52).

Synteny and comparative genomics were performed in R using custom scripts where one representative genome per SEA-PHAGES subcluster (D29 for A2) was selected, gene neighbourhoods of ±5 ORFs around gp53 were extracted and drawn relative to the gp53 midpoint, and the six most conserved neighbouring phams were coloured, with gp52 (pham 187767) and gp53.1 (phams 85069/84872/113591/173799) coloured explicitly. For every Cluster A genome encoding both genes, the gp53–gp52 archi-tecture (fused, overlapping, 0-nt spacer, or separated by an intergenic spacer) was tabulated, and for every observed spacer length the pham of the first intervening ORF was taken from the Phamerator gene table; cassette phams were confirmed by direct hmmscan of the intervening ORFs.

For co-evolution analysis, gp53 and gp52 homologs were paired by genome identi-fier, retaining the 644 genomes that encode a standalone (non-fused) copy of each.

Paired protein sets were aligned independently with MAFFT (--auto) and trimmed with trimAl (-automated1). Maximum-likelihood trees were inferred for each set with IQ-TREE v3.1.2 (ModelFinder; 1,000 ultrafast-bootstrap replicates). Patristic (cophe-netic) distance matrices were obtained from both trees (ape::cophenetic.phylo), and MirrorTree [60] congruence was quantified as the Pearson correlation between the upper-triangle entries of the gp53 and gp52 matrices (*r* = 0.61). Significance was assessed by permutation of gp52 tip labels (5,000 permutations) using an empirical two-sided *p*-value, *p* = (1 + #{|*r*_perm_| ≥ |*r*_obs_|})*/*(*N*_perm_ + 1) (*p* ≈ 2 × 10*^−^*^4^); the same correlation computed directly on pairwise amino-acid distances (ape::dist.aa, pairwise deletion) gave *r* = 0.86. Topological congruence was measured as the Robinson–Foulds distance (ape::dist.topo, PH85) normalised as RF*/*(2*n*−6), where *n* is the number of shared tips (0.92, *n* = 644). To assess co-occurrence independently of tree shape, gp53/gp52 presence–absence was computed across the full genome panel and association tested with Fisher’s exact test. Unless otherwise specified, tools were run with default parameters and random seeds were fixed for permutation analyses.

## Acknowledgements

We thank Graham Hatfull (University of Pittsburgh, PA, USA) for providing the D29 phage used in this study. We thank the members of the AF Lab for their valuable discussions and insights. AF, SG, DK and GIP are supported by the SciLifeLab and Wallenberg Data-Driven Life Science Program (KAW 2020.0239) and the Swedish Research Council (2023-01970 VR). DP is supported by the KI-SciLifeLab Clinical Translational Postdoc Program funded by the Strategic Research Area (Strategiskt Forskningsområde, SFO) for Molecular Life Science at Karolinska Institutet. to AF This work was supported by SNF RU Institute for Global Infectious Disease Research at The Rockefeller University (PDBO. and EAC.). AE and SND are funded by Veten-skapsrådet, Grant No. 2021-03979, the Knut and Alice Wallenberg Foundation, Grant No. 2022.0032, and the Swedish E-science Research Center. The computations and data handling were enabled by the supercomputing resource Berzelius, provided by the National Supercomputer Center at Linköping University, the Knut and Alice Wallen-berg Foundation, and SNIC, grant numbers SNIC 2021/5-297 and Berzelius-2021-29. The funders had no role in study design, data collection and analysis, decision to publish or preparation of the manuscript.

## Author contributions

D.R.P.: Methodology, Investigation, Formal Analysis, Writing — Review & Editing. S.G.: Methodology, Investigation, Formal Analysis, Writing — Review & Editing. A.D.: Methodology, Investigation, Formal Analysis, Writing — Review & Editing. D.P.O.: Methodology, Investigation, Formal Analysis, Writing — Review & Editing.

S.N.D. : Methodology, Formal Analysis. G.I.P.: Investigation. D.K.: Investigation. F.A.R.: Investigation. A.E.: Resources, Methodology, Supervision. M.M.: Resources, Methodology, Supervision. E.A.C.: Resources, Supervision, Methodology, Formal Analysis, Writing — Review & Editing, Funding Acquisition. A.F.: Resources, Fund-ing Acquisition, Conceptualization, Methodology, Supervision, Writing — Review & Editing.

## Supplementary Figures

**Fig. S1.**
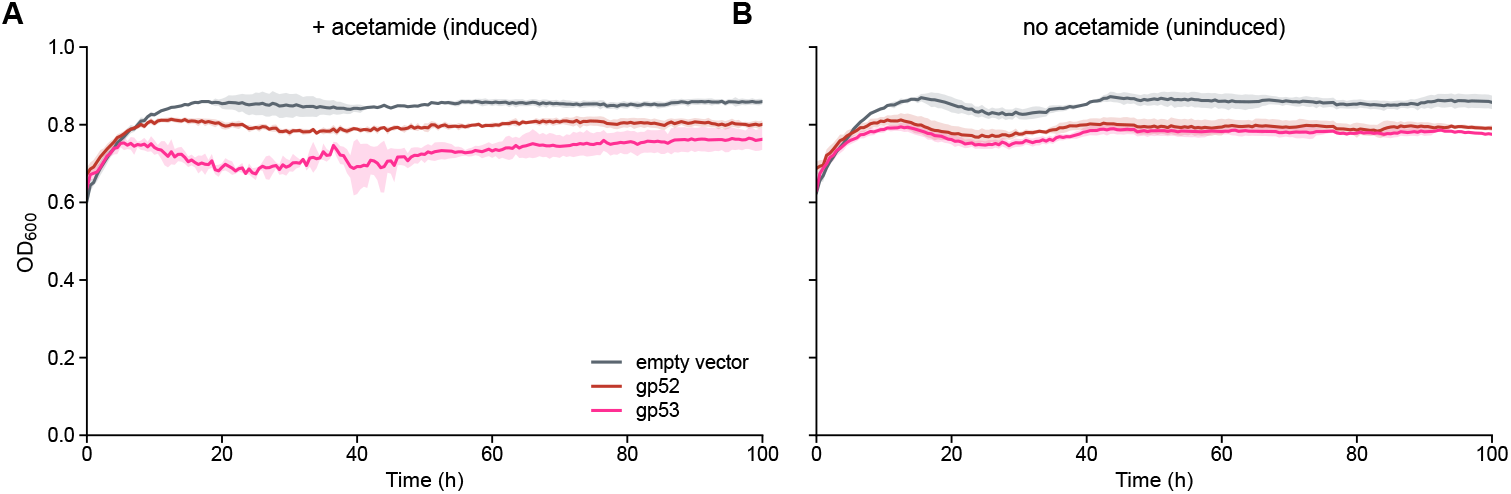
Growth of *M. smegmatis* expressing gp53 or gp52. Growth of *M. smegmatis* carrying an acetamide-inducible Strep-tagged gp53 or gp52 construct, or the empty vector, followed as OD_600_ measurements over 100 h. In both panels the x-axis shows time after inoculation in hours and the y-axis shows OD_600_; the two panels share the same y-axis. Colour represents the strain: grey, empty vector; red, gp52; magenta, gp53. Lines show the mean of *n* = 3 wells and shaded bands the SD. **A.** Cultures induced with acetamide. **B.** The same strains without inducer.

**Fig. S2.**
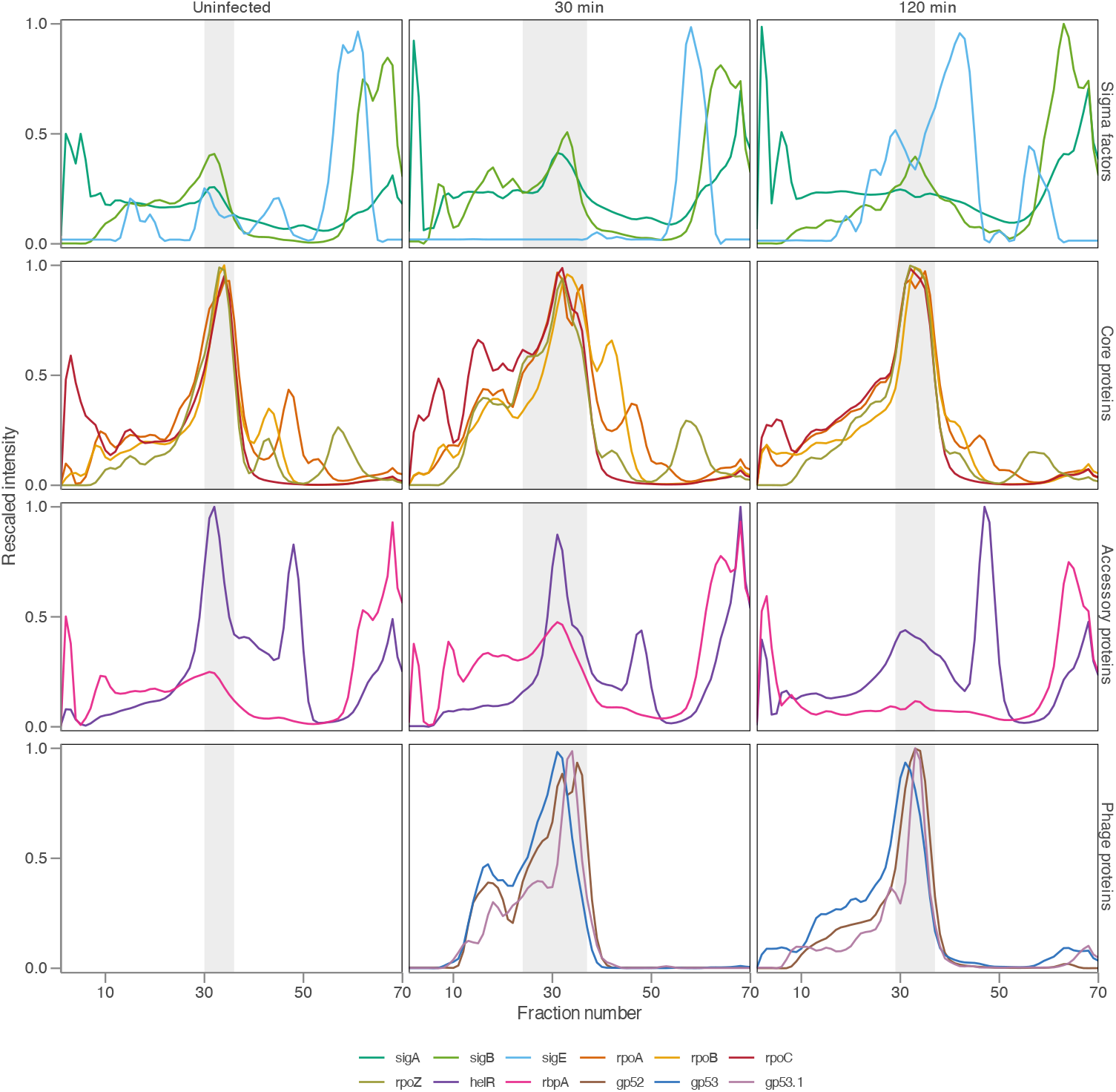
Co-elution profile for RNA polymerase components and phage proteins. Indi-vidual RNA polymerase-associated fractionation traces grouped by protein class. Traces show mean rescaled intensity across biological replicates for each protein, with one line per protein. Rows show sigma factors, core RNA polymerase proteins, accessory proteins, and phage proteins (gp52, gp53, gp53.1); columns show uninfected, 30 min, and 120 min conditions. Light grey shading marks the core RNA polymerase peak region, defined from the half-maximal window of the averaged core-protein signal in each condition. The phage-protein panel is left empty for the uninfected condition. X-axis: fraction number; y-axis: rescaled intensity.

**Fig. S3.**
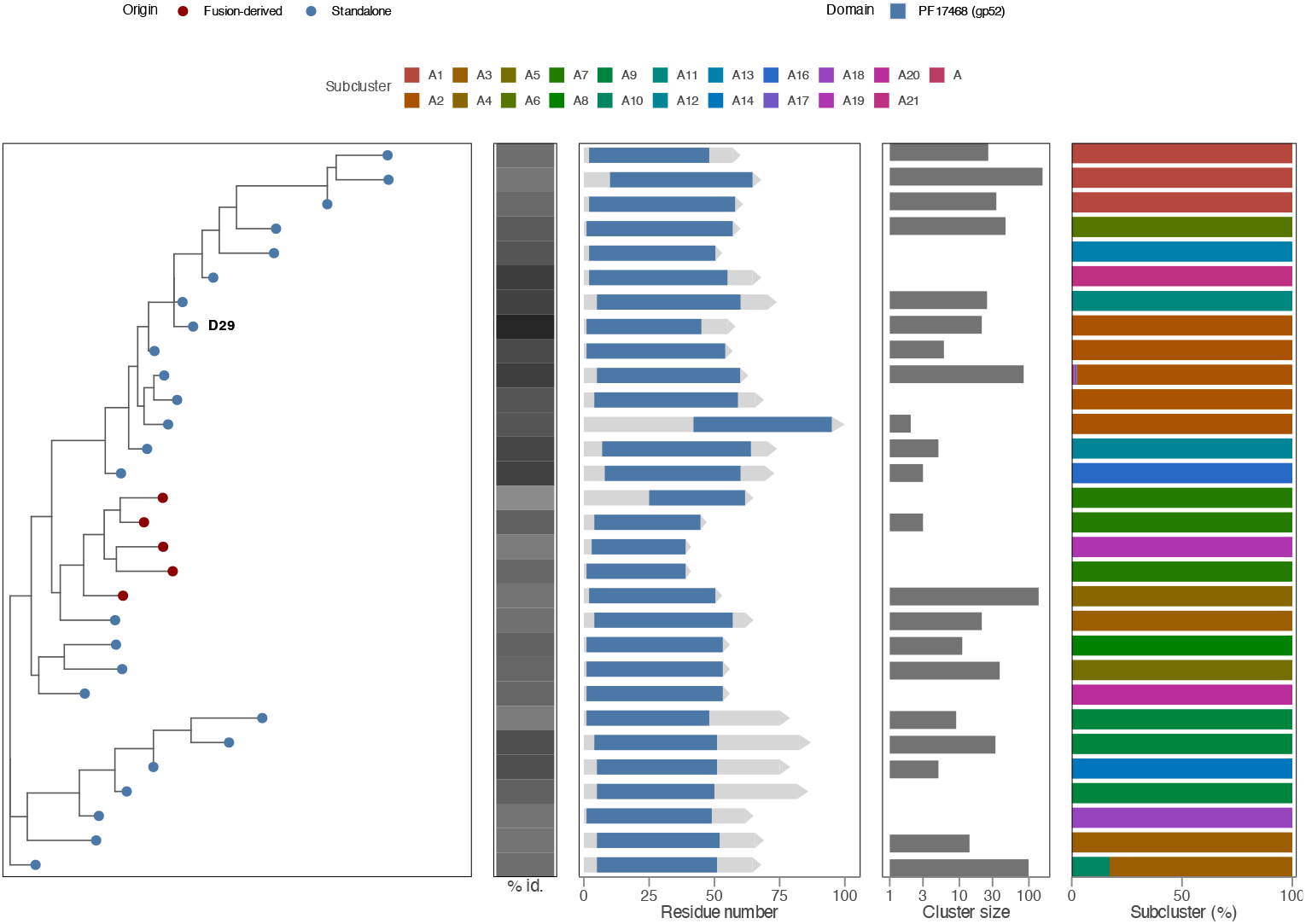
Evolutionary relationships, domain architecture and conservation of gp52 across phage subclusters. Rectangular maximum-likelihood phylogenetic tree of 30 gp52 sequence-cluster representatives (MMseqs2, 80% identity clustering of the gp52 union set), with the representative carrying D29 gp52 labeled and four row-aligned tracks. Tip dots are coloured by the origin of the gp52 sequence: Standalone (drawn from a separate gp52 gene) or Fusion-derived (the PF17468 region truncated out of a longer fused gp53–gp52 protein). *% id.*, grayscale heatmap of percent amino acid identity to D29 gp52. *Residue number*, to-scale gene-arrow schematic of each representative’s protein length showing the PF17468 (GP52) domain block; Fusion-derived entries were already truncated to their PF17468 span when the gp52 union was built, so no HTH portion is shown. *Cluster size*, the number of collapsed duplicate sequences each representative stands for (log scale). *Subcluster (%)*, the SEA-PHAGES subcluster composition of each representative’s cluster members. This panel is the gp52 analogue of Fig. 3A and was built from an independently inferred gp52 tree.

**Fig. S4.**
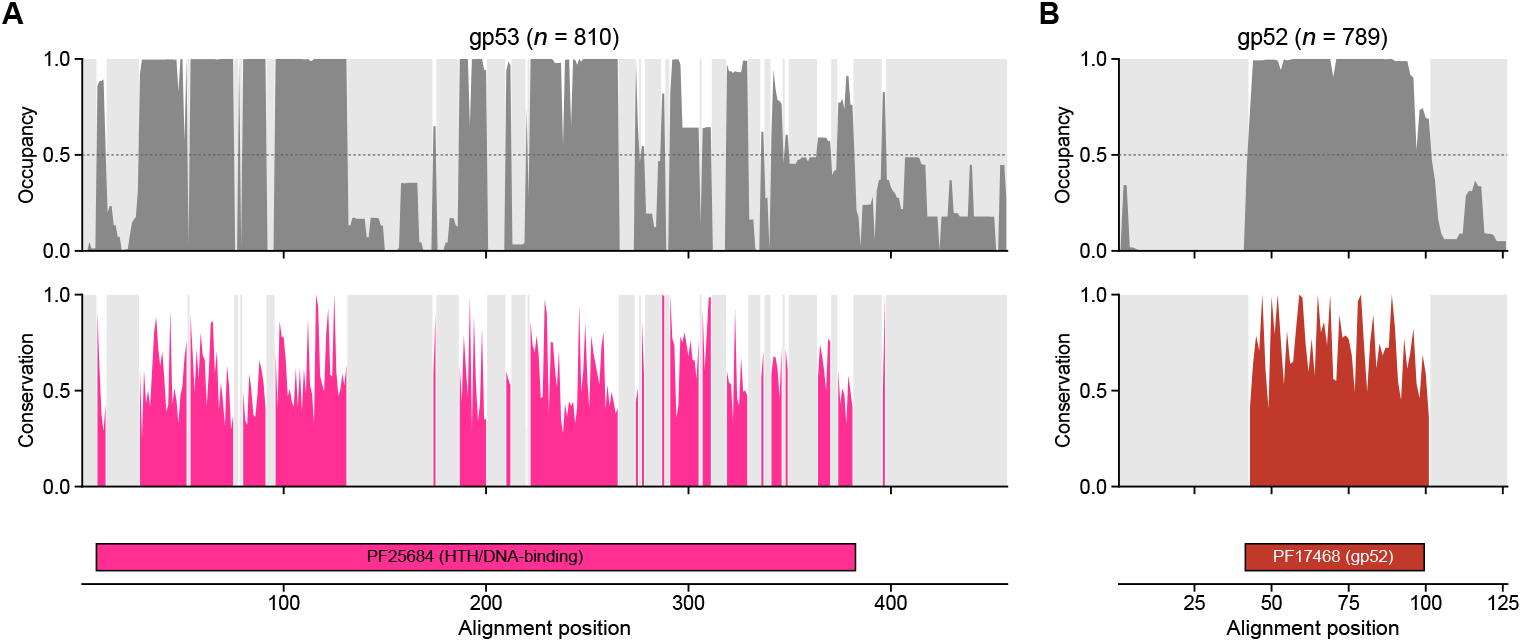
gp53/gp52 conservation and inferred domain architecture. In both panels the x-axis shows the position in the multiple-sequence alignment. The top track shows per-column occupancy, the fraction of sequences carrying a residue rather than a gap, and the middle track shows per-column conservation; both y-axes run from 0 to 1, and the dashed line marks 50% occupancy. Columns below 50% occupancy are shaded grey and masked in the conservation track. The bottom track marks the extent of the Pfam domain. Colour represents the protein: magenta, gp53; red, gp52. **A.** gp53-family proteins (*n* = 810), with the PF25684 helix-turn-helix/DNA-binding region marked. **B.** gp52-family proteins (*n* = 789), with the PF17468 gp52 region marked.

